# Analysis of stochastic timing of intracellular events with gene switching

**DOI:** 10.1101/710442

**Authors:** Khem Raj Ghusinga, Abhyudai Singh

## Abstract

An important step in execution of several cellular processes is accumulation of a regulatory protein up to a specific threshold level. Since production of a protein is inherently stochastic, the time at which its level crosses a threshold exhibits cell-to-cell variation. A problem of interest is to characterize how the statistics of event timing is affected by various steps of protein expression. Our previous work studied this problem by considering a gene expression model where gene was always active. Here we extend our analysis to a scenario where gene stochastically switches between active and inactive states. We formulate event timing as the first-passage time for a protein’s level to cross a threshold and investigate how the rates of gene activation/inactivation affect the distribution and moments of the first-passage time. Our results show that both the time-scale of gene switching with respect to the protein degradation rate as well as the ratio of the gene inactivation to gene activation rates are important parameters in shaping the event-timing distribution.

## I. INTRODUCTION

In the ever-changing cellular environment, a cell has to actively sense and respond to both internal and external cues [1], [2]. The environment is represented via transcription factors. A change in the environment leads to localization of appropriate transcription factors which subsequently activate expression of target genes [3]–[5]. The temporal aspect of the target gene activity plays important role in cellular function as cells often couple important decisions or events to accumulation of specific regulatory proteins up to critical thresholds. Many examples of such events appear in context of development [6]–[11], cell-cycle control [12]–[18], cell differentiation [19]–[23], sporulation [24], [25], apoptosis [26]–[28], lysis of infected bacterial cells [29]–[31], etc.

As the protein levels are subject to molecular fluctuations due to inherent noise in gene expression [32]–[42], the timing of an event that triggers at a critical threshold is expected to exhibit cell-to-cell variation. Indeed, recent single-cell experiments have shown considerable cell-to-cell variability in timing of cellular events [29], [43].

Our recent work has characterized the stochasticity in the timing of events originating from the inherent gene expression noise by formulating event timing as a first-passage time problem [44], [45]. The models of gene expression considered therein however only accounted for transcription (synthesis of mRNA from gene) and translation (synthesis of protein from mRNA) while assuming that the gene was always active. However, single-cell measurements have revealed that genes are not always active. Instead, they switch between active and inactive states [46]–[54].

Here we explore the effects of gene switching on the first-passage time statistics. We model the gene dynamics as a telegraph process that switches between ON (active) and OFF (inactive) states at exponentially distributed times. We specifically examine the effect of the average time period of the pulsing, and the average duty cycle (average fraction of time for which the signal stays in ON state) on the FPT statistics. In addition to gene switching, our model considers production of mRNAs when gene is in the ON state. Regardless of the state of the gene, each mRNA can synthesize proteins before degrading. For analytical tractability, we assume that mRNA half-life is small as compared to that of the protein of our interest. Therefore, we can ignore its dynamics and assume that each transcription event produces protein molecules in bursts that follow geometric distribution [55]–[57].

Our results show that two important parameters that affect the shape of the distribution as well as the moments of event timing are: the time-scale of switching with respect to the protein degradation rate, and the ratio of gene activation rate to gene inactivation rate. The effect of time-scale is more prominent when the gene activation rate is smaller than or comparable to the gene inactivation rate.

The paper is organized as follows. Section II provides detailed description of our model and then discusses the calculations for first-passage time for the model considered herein. Section III explores the effects of gene switching parameters on the first-passage time statistics, and finally Section IV concludes the paper.

## II. STOCHASTIC DESCRIPTION OF EVENT TIMING

In this section, we describe a model of gene expression and use it to compute statistics of event timing.

### A. Model Formulation

Let *g*(*t*) ∈ {0, 1} denote the state of the gene such that *g*(*t*) = 0 implies that the gene is inactive (OFF) and *g*(*t*) = 1 represents that the gene is active (ON). The stochastic switching of the gene between these two states is modeled as a telegraph process

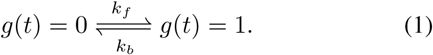

Once the gene is in the active state, it produces mRNAs with a rate *k*. In other words, the mRNA synthesis rate is given by *kg*(*t*). Each mRNA molecule produces proteins with a rate *k*_*p*_ and it degrades with rate *γ*_*m*_. The protein degradation rate is denoted by *γ*_*p*_ (Fig. 1). Both mRNA transcripts and the protein molecules degradation events are considered to be single step processes.

**Fig. 1.**
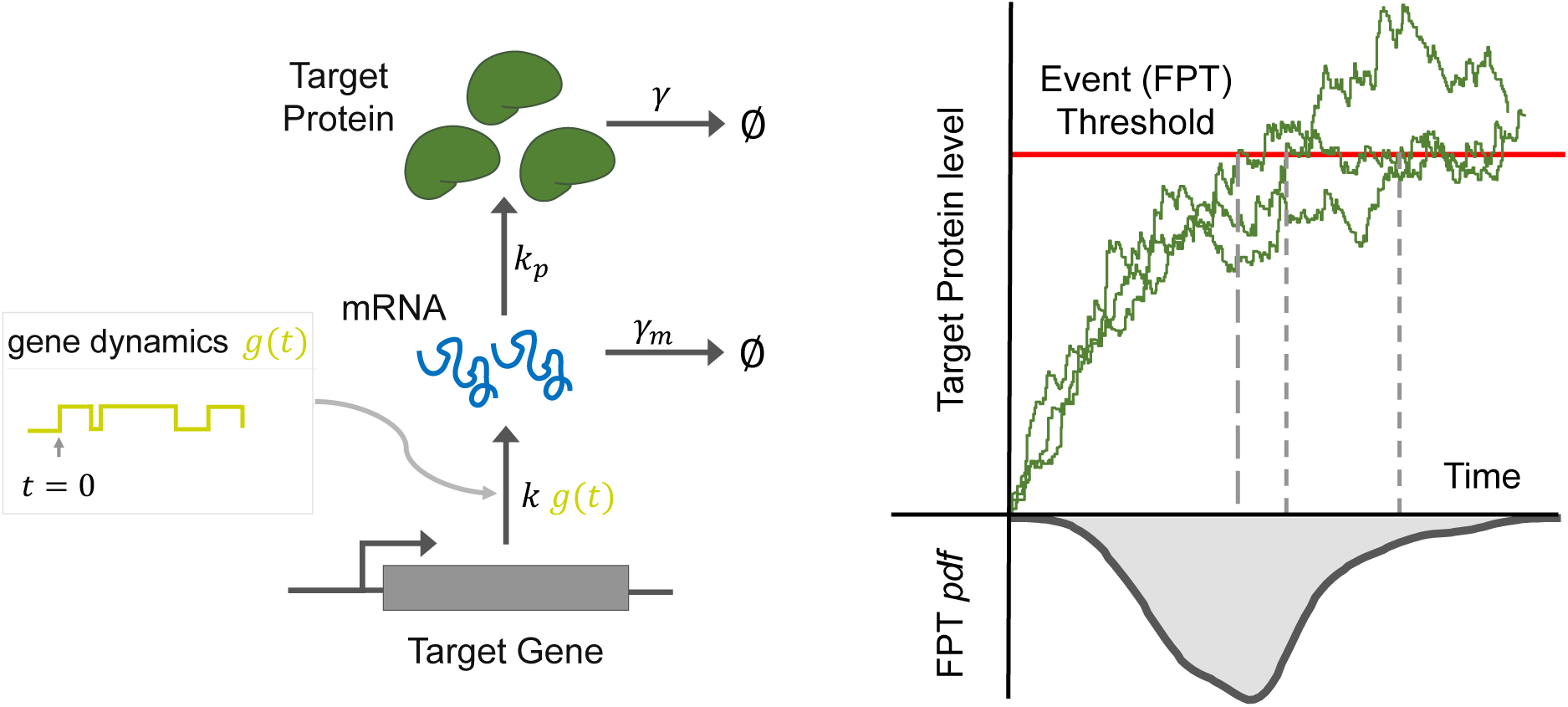
Event timing as a first-passage time problem. Left: A sketch of the gene expression model that includes production and degradation of mRNA/protein molecules. The gene dynamics is represented by *g*(*t*). Right: An event of interest is triggered when the protein level reaches a critical event threshold for the first time. Each protein trajectory is representative of protein level over time inside individual cells. As a consequence of stochastic expression of the protein, the threshold is attained at different times in different cells. The corresponding time at which the event happens denotes the first-passage time/event time.

For analytical tractability, we assume that mRNA half-life is small as compared to that of the protein of our interest, i.e., *γ*_*m*_ ≫ *γ*_*p*_. Therefore, we can ignore its dynamics and assume that each transcription event produces protein molecules in bursts that follow geometric distribution with mean burst size *b* = *k*_*p*_*/γ*_*m*_ [55]–[57]. More precisely, let *x*(*t*) denote the protein level in a cell at time *t* then the probabilities of gene switching, protein production, and degradation events taking place in an infinitesimal time interval (*t, t* + *dt*) as

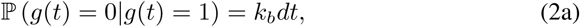

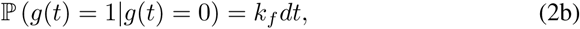

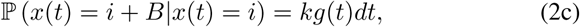

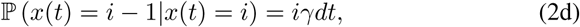

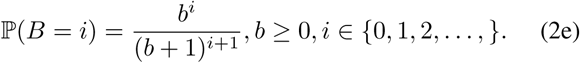

Here, ℙ is notation for probability and *B* denotes the burst size which follows a geometric distribution with mean *b*.

Having defined the model, we can now characterize the statistics of the time at which protein level crosses a specific threshold.

### B. Event Timing Distribution

We are interested in computing distribution of the first time at which *x*(*t*) crosses a threshold *X* (Fig. 1). The time to this event can be conveniently formulated as first-passage time (FPT) defined as

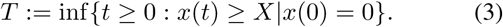

The pdf of *T* can be computed by constructing an auxiliary process *x*_*aux*_(*t*) on the state-space {0, 1,…, *X*}. The initial conditions as well as the probabilities of occurrences for this process are exactly same as *x*(*t*) in (2), except that here *X* is an absorbing state, i.e., once the process reaches the state *X*, it remains there forever. In terms of the first-passage time, both processes *x*(*t*), and *x*_*aux*_(*t*) are identical.

The probability that the state *X* is reached by *x*_*aux*_(*t*) in the time-interval (*t, t* + *dt*) is given by

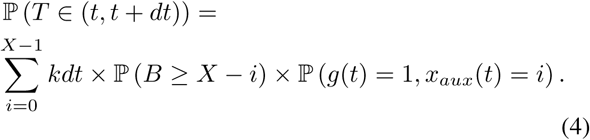

Denoting the pdf of *T* by *f*_*T*_ (*t*), using the fact that Probability (*B* ≥*X* − *i*) = (*b/b* + 1)^*X-i*^, and denoting *P*_1,*i*_(*t*):= P (*g*(*t*) = 1, *x*_*aux*_(*t*) = *i*), we get

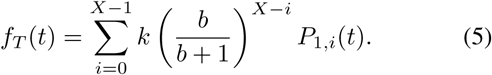

The terms *P*_*i*_(*t*) can be determined from the Chemical Master Equation (CME) which can be written as a system of differential equations

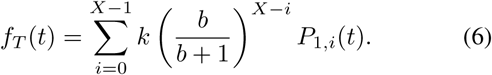

The terms *P*_1,*i*_(*t*) can be computed from the master equation for the auxiliary process that can be written as

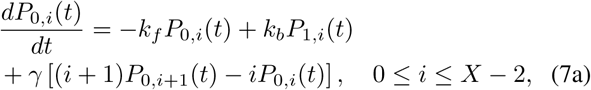

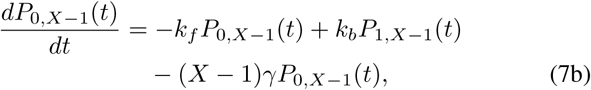

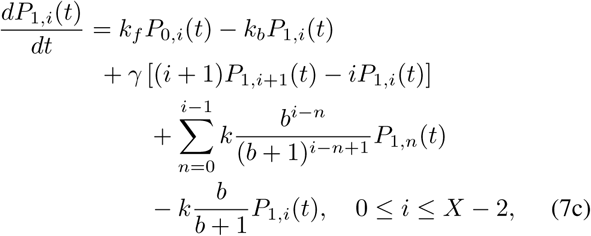

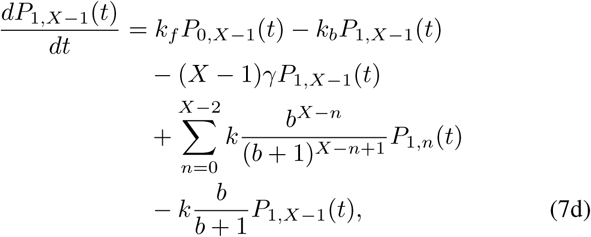

where we have used the following notation

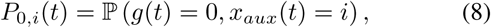

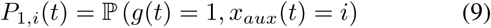

[58]–[61]. Stacking the probabilities in a single vector

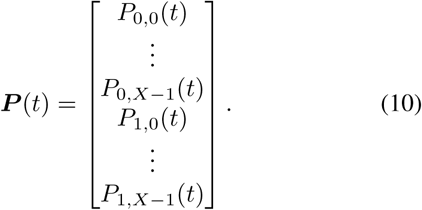

leads to a convenient form of the CME as the following linear time invariant dynamical system

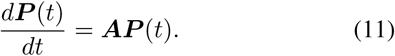

The matrix ***A*** in the above equation can be written as

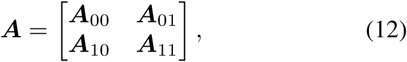

where (*i, j*) − *th* elements of the *X* × *X* block matrices are given by

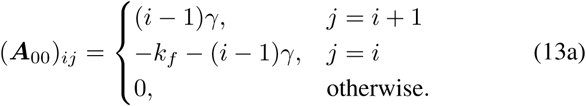

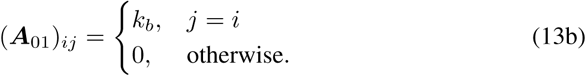

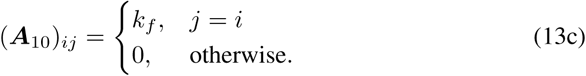

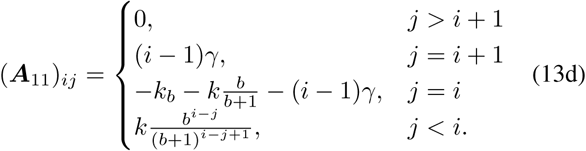

where *i, j* ∈ {1,…, *X*}.

Using the solution of the linear time invariant system of equations in (10), the pdf of *T* can be written as

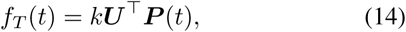

where ***U*** represents the following 2*X*-dimensional vector

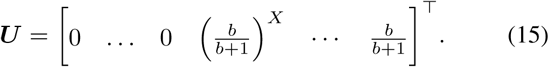

The solution ***P*** (*t*) can be analytically computed as

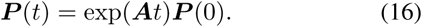

Here the initial condition is taken as ***P*** (0) = 0… 0 1 0… 0]^T^ which corresponds to *g*(0) = 1, *x*_*aux*_(0) = *x*(0) = 0 with probability one. As a result, the pdf of *T* takes the form

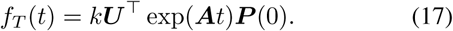

We can now use (17) to compute the moments of *T*. In particular, we study the first two moments in this work.

### C. Moments of Event Timing

We can use (17) to compute a *m*^*th*^ order moment as

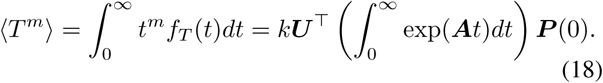

Though we do not provide a formal proof here, the matrix ***A*** has the property that its diagonal elements are negative and dominant (i.e., sum of absolute values of all other elements in a column is less than absolute value of the diagonal element). Thus, we can use the result from [62, Thm. 1, pp 48–49] to argue that ***A*** is full-rank and eigenvalues with negative real parts. Therefore, the above integral simplifies t o [45]

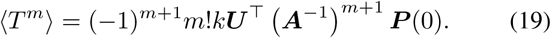

To compute ***A***^*-*1^, we now use the standard formulas for inverse of a block diagonal matrix as [63]

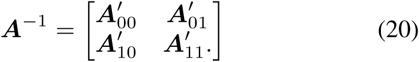

where

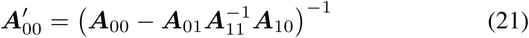

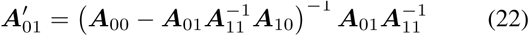

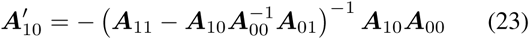

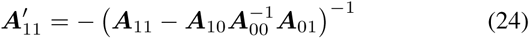

Typically, there are multiple ways to write the inverse, depending upon which of the block matrices are non-singular. Due to sheer complexity of the expressions, we do not provide explicit form of ***A***^*-*1^. At any rate, we can numerically invert these matrices and obtain ***A***^*-*1^ to compute the moments.

So far, we have found an expression for the distribution and moments of *T* in terms of the model parameters. In the next section, we explore the effect of model parameters on these.

## III. EFFECT OF MODEL PARAMETERS

We begin by exploring the effect of gene switching parameters on the distribution of *T* given by (17). To keep the analysis relative to the protein time-scale, we set *γ*_*p*_ = 1. In Fig. 2, we plot the distribution for various values of *k*_*f*_ and *k*_*b*_. Since the time-scale of gene turning off is set by 1*/k*_*b*_, we look at its values that are orders of magnitude different. Furthermore, the average time for which the gene remains active is given by 1*/*(1 + *k*_*b*_*/k*_*f*_). Therefore, we look at various values of *k*_*f*_ that span a couple of orders of magnitude of *k*_*b*_. We note that when *k*_*b*_ is small as compared to *γ*, i.e., the switching occurs at a slow rate, then the ratio of *k*_*f*_ to *k*_*b*_ doesn’t affect the distribution much (blue curves). On the other hand, if *k*_*b*_ is much higher than *γ*, then the ratio greatly impacts the distribution shape (magenta curves). In particular, when *k*_*b*_ is higher than *k*_*f*_, the distribution has a heavy-tail (top, right). Likewise, when *k*_*f*_ is much larger than *k*_*b*_, then the relative value of *k*_*b*_ with respect to *γ* doesn’t affect the distribution much (bottom row). The value of *k*_*b*_ starts to impact the distribution when *k*_*f*_ is either similar or much smaller than *k*_*b*_ (middle and top rows).

**Fig. 2.**
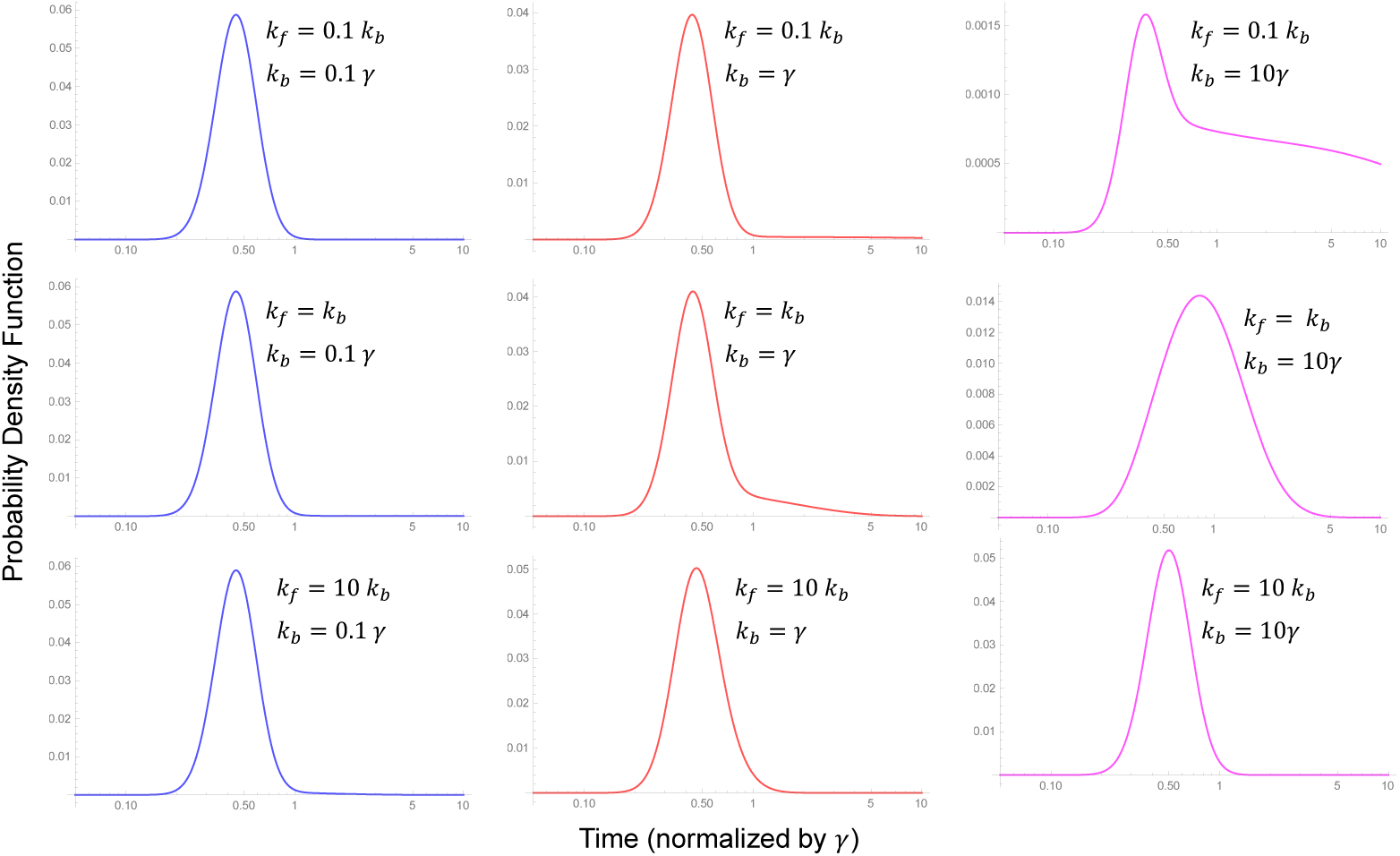
Effect of gene switching parameters on the probability density function of T in (17). We have chosen *γ* = 1, so as to normalize all parameters with respect to the protein degradation rate. The values of *k*_*f*_ and *k*_*b*_ for each plot are mentioned therein. Rest of the parameter values are chosen as *k* = 50, and *X* = 20, and *b* = 1.

In our previous work [64], we noted that the relative positions of the study-state protein level with respect to the event threshold can affect the noise in event timing in non-trivial ways. In this model, the steady-state protein level is given by

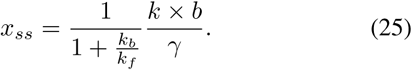

Indeed when the distribution has a long-tail (as in top row, right), *x*_*ss*_ = 50*/*11 which is much less than the event threshold *X* = 20. Therefore, the relative positions of *x*_*ss*_ and *X* seem to matter in this case as well.

We can also use the expression of moments in (19) to look at the effect of moments. To illustrate this, we look at the effect of duty cycle *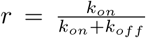*, i.e., fraction of time the gene is in ON state on the moments of event timing. Interestingly, while the mean time expectedly decreases with increase in the duty cycle, the noise (quantified by coefficient of variation squared) shows a non-monotonic behavior (Fig. 3). Note that while the decrease in noise can be explained by the relative positions of the event threshold and *x*_*ss*_, the earlier increase in the noise cannot be explained this way since *x*_*ss*_ increases with duty-cycle.

**Fig. 3.**
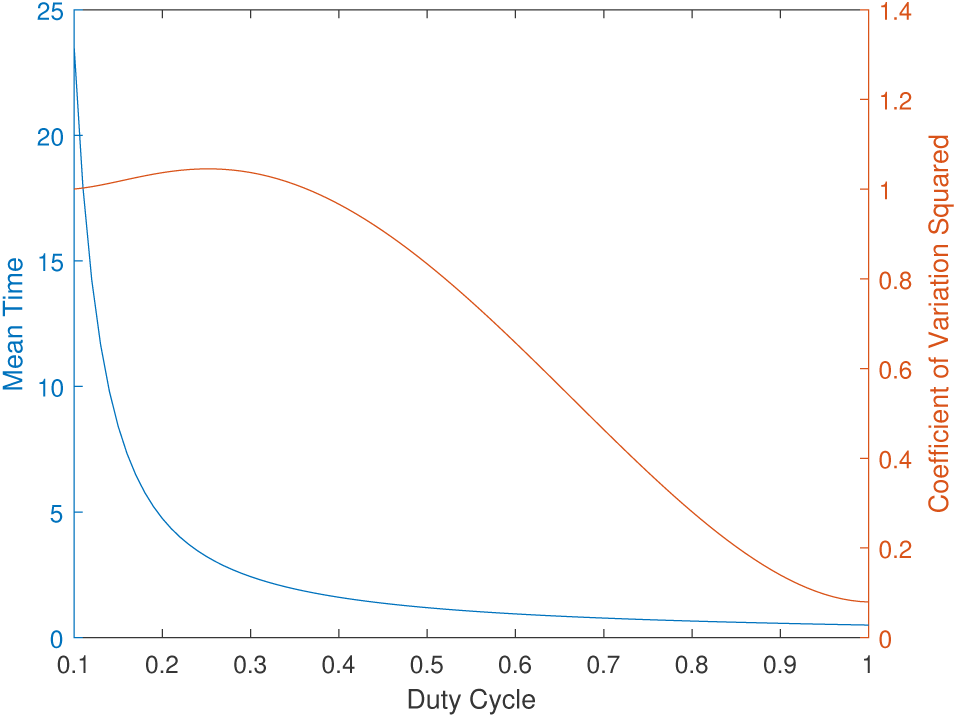
Effect of gene switching parameters on the moments of T in (19). We have chosen *γ* = 1, so as to normalize all parameters with respect to the protein degradation rate. Rest of the parameter values are chosen as *k* = 50, and *X* = 20, and *b* = 1.

## IV. CONCLUSIONS

In this paper, we studied the effect of gene switching on the distribution and statistics of timing of an event that triggers upon attainment of a specific protein level. We showed that both the ratio of switching rates as well as the time-scale of switching as compared to the protein degradation rate affect the statistics. Future work will explore effect of feedback strategies on event timing, wherein the feedback affects the rate of activation or inactivation.

